# Effect of mechanical properties of monofilament twines on the catch efficiency of biodegradable gillnets

**DOI:** 10.1101/2020.05.22.110379

**Authors:** Eduardo Grimaldo, Bent Herrmann, Nadine Jacques, Jørgen Vollstad, Biao Su

## Abstract

Gillnets made of the biodegradable resin polybutylene succinate co-adipate-co-terephthalate (PBSAT) were tested under commercial fishing conditions to compare their fishing performance with that of conventional nylon polyamide (PA) gillnets. Both types of gillnets were made of 0.55 mm Ø monofilaments. However, since the biodegradable nets are weaker than nylon PA nets when using the same monofilament diameter, we also used biodegradable nets made of 0.60 mm Ø monofilament that had a similar tensile strength to the 0.55 mm Ø nylon PA nets. The relative catch efficiency of the different gillnet types was evaluated over the 2018 autumn fishing season for saithe and cod in northern Norway. For cod, both biodegradable gillnets (0.55 and 0.60 mm) had a significantly lower catch efficiency compared to the traditional nylon PA net (0.55 mm) with estimated catch efficiencies of 62.38% (CI: 50.55–74.04) and 54.96% (CI: 35.42–73.52) compared with the nylon PA net, respectively. Similarly for saithe, both biodegradable gillnets (0.55 and 0.60 mm) had a lower estimated catch efficiency compared to the traditional nylon PA net (0.55 mm) with estimated catch efficiencies of 83.40% (71.34–94.86) and 83.87% (66.36–104.92), compared with the nylon PA net, respectively. Tensile strength does not explain the differences in catch efficiency between the two gillnet types, since increasing the twine diameter of the biodegradable gillnets (to match the strength of nylon PA gillnets) did not yield similar catch efficiencies. However, the elasticity and stiffness of the materials may be responsible for the differences in catch efficiency between the nylon PA and biodegradable gillnets.

## Introduction

Globally, gillnets are among the most commonly used fishing gears in developing and industrialized countries [1]. In Norway, 26% and 16% of the total national allowable quota for Northeast Atlantic cod (*Gadus morhua*) and saithe (*Pollachius virens*), which in 2019 was 385.000 and 203.368 tonnes respectively, were caught with gillnets [2]. The Norwegian coastal fleet (with vessels shorter than 28 m) is responsible for approximately 99% of the gillnet landing of Northeast Atlantic cod. In 2019, the coastal fleet consisted of 5978 vessels, with 81% of them being smaller than 14.9 m [3]. Despite the importance of the gillnet fishery, large numbers of gillnets are lost every year, causing environmental problems such as ghost fishing and marine litter. Deshpande et al. [4] provided annual loss rates of the six types of fishing gears used in Norwegian waters upon deployment, and gillnets were the primary source of derelict gear. Although fisheries authorities lack a complete overview of the amount of lost or derelict gillnets, estimates from the Norwegian Environment Agency [5] suggest that 13,941 gillnets are lost each year.

The impacts of derelict gillnets include continued catching of target and non-target species (commonly known as ghost fishing), alterations to the benthic environment, marine plastic pollution, navigational hazards, beach debris/litter, introduction of synthetic material into the marine food web, costs related to clean-up operations, and impacts on business activities [6]. The impact of derelict gillnets on the environment has been exacerbated by the introduction of non-biodegradable materials, primarily plastics, which are generally more persistent in the environment than natural materials. With reference to the principles for fisheries resource management (the Gordon-Schaefer model) [7], ghost fishing also represents an unregistered amount of fishing mortality, which undermines the use of the population analysis models for maximum sustainable yield management and the ecosystem management approach. There have been extensive efforts to assess the magnitude of derelict gillnets [8, 9], and in the last decade many studies have focussed on developing methods to reduce the effects of derelict gear. Some specific measures to address the problem include gear marking, onshore collection/reception and/or payment for old/retrieved gear, reduced fishing effort, use of biodegradable nets, and gear recovery programs for gear disposal and recycling [9].

Norway is one of the countries that has a program to systematically retrieve lost gears from areas with the highest fishing intensity. Between 1983 and 2017, the Norwegian Directorate of Fisheries retrieved 20,450 lost gillnets and a large amount of other fishing gear (e.g., ropes, pots, trawls), which contained variable amounts of marine resources that had been caught in the lost gear (ghost fishing). In 2017, just 815 of the 13,941 gillnets that were reported lost were retrieved [5, 10]. Due to the low recovery rate of lost fishing gears and the low on-land disposal rate of plastics from the fishing industry, recent research has focused on developing biodegradable plastic materials for fishing gear, i.e. gillnets, to try to reduce the negative effects of derelict fishing gear.

Biodegradable plastic is a plastic that maintains the same properties as a conventional plastic during use, but that can be completely degraded by naturally occurring microorganisms such as bacteria, fungi, and algae when disposed of in the environment [11]. The most investigated biodegradable plastics in fishing equipment and marine applications, i.e., aquaculture, are polybutylene succinate (PBS), polybutylene adipate co-terephthalate (PBAT), and polybutylene succinate co-adipate-co-terephthalate (PBSAT) [12-20]. Commercial fishing products made of these materials are available in some countries, such as South Korea. Other biodegradable plastics of interest include polylactic acid (PLA), polycaprolactone (PCL), polybutylene succinate adipate (PBSA), polyhydroxyalkanoates (PHAs) (e.g., poly(3-hydroxybutyrate) [P(3HB)] and poly(3-hydroxybutyrate-co-3-hydroxyvalerate) [P(3HB-co-3HV)], and combinations of PHAs. Various microorganisms are known to degrade biodegradable plastics at different rates, for example, the microorganisms present in the Arctic have a high capacity for biodegradation [21]. Additionally, there are reports that the degradation of PCL and PHB/V fibres occurs at a faster rate than that of PBS fibres in deep seawater [22]. However, PBSA may be degraded by several microorganism types compared to PBS [21]. Biodegradable fishing nets have thermal, mechanical, and physical properties that are comparable to those of traditional products made of nylon polyamide (PA), polyester (PES), polyethylene (PE), and polypropylene (PP) [12, 13, 17].

Biodegradable fishing gears have been studied in South Korea and Norway as an alternative to reduce the negative impact of derelict gear on the marine environment. In South Korea, these gears have been tested in 13 different fisheries, including gillnetting and potting for roundfish, flatfish, shrimps, octopus, crabs, and eels [12-17, 23-25]. The results showed that in some cases the fishing efficiency of these gears is similar to that of gears made of PA, PE, and PP. In Norway, biodegradable gillnets have shown a consistently lower catch efficiency than nylon PA gillnets, and this difference has been mainly attributed to the fact that biodegradable gillnets are made with 11-16% weaker monofilaments than nylon PA monofilaments of the same diameter [18-20, 26]. The aim of the present study is to assess the effect of twine thickness tensile strength on the catch efficiency of biodegradable gillnets. Our main hypothesis is that by increasing the monofilament diameter of the biodegradable gillnets, to match the tensile strength of nylon PA monofilaments, the catch efficiency of the biodegradable gillnets will be improved and yield a similar catch efficiency to nylon PA gillnets. We designed the fishing experiments to answer the following research questions:

i. Can biodegradable and nylon PA gillnets made of monofilaments with similar tensile strength (although different monofilament diameter) yield similar catch efficiencies?
ii. Is tensile strength the mechanical property responsible for the difference in catch efficiency between biodegradable and nylon PA gillnets?
iii. Is catch efficiency positively or negatively correlated to monofilament diameter in biodegradable gillnets?

## Materials and methods

### Ethics Statement

This study did not involve endangered or protected species. Experimental fishing was conducted on board a commercial fishing vessel and no permit was required to conduct the study on board. No information on animal welfare, or on steps taken to mitigate fish suffering and methods of sacrifice is provided, since the animals were not exposed to any additional stress other than that involved in commercial fishing practices.

### Experimental setup

Sea trials were conducted on board the coastal gillnet vessel “MS Karoline” (10.9 m total length) throughout October and December 2018. The fishing grounds chosen for the sea trials were located off the coast of Troms (Northern Norway) between 70°21’–70°22’N and 19°39’–19°42’E, which is a common fishing area for coastal vessels from Troms targetting cod and saithe.

A 130 mm nominal mesh opening was used for both types of gillnets, with monofilament twine thickness of 0.55 and 0.60 mm in the biodegradable gillnets and 0.55 mm in the nylon PA gillnets. Since the biodegradable monofilament is considered to be approximately 10% weaker than nylon PA monofilament (at equal monofilament thickness), the monofilament thickness was increased from 0.55mm to 0.60 mm to compensate for the difference in tensile strength.

Two sets of gillnets were used in the experiments. Each set consisted of 16 gillnets, with eight biodegradable gillnets (B) and eight nylon PA gillnets (N). The gillnets were arranged in such a way that they provided information for paired comparison, nylon PA versus biodegradable gillnet, accounting for spatial and temporal variation in the availability of cod. With individual sets being the basic unit for the paired analysis [19], it was important that the biodegradable and nylon PA gillnets were approximately exposed to the same spatial variability in cod availability within each gillnet set. This could in principle be achieved by alternating between the two types of nets after each net sheet as follows: B-N-B-N-B-N-B-N-B-N-B-N-B-N-B-N. However, for ease of on board recording of fish in relation to the type of net in which it was caught, the alternation in net types was only applied after every second net sheet. Therefore, to make conditions as equal as possible between net types, set 1 was arranged as N-BB-NN-BB-NN-BB-NN-BB-N and set 2 as B-NN-BB-NN-BB-NN-BB-NN-B. Actual measurements of the mesh openings (four rows of 20 meshes each) were taken with a Vernier calliper without applying tension to the meshes, which showed that the mean mesh openings of 0.55mm nylon PA gillnets and 0.55mm and 0.60mm biodegradable gillnets were 131.6 ± 0.72 mm, 131.5 ± 1.0 mm and 132.5 ± 0.8 mm, respectively.

### Data analysis

#### Modelling catch efficiency

We used the statistical analysis software SELNET [27, 28] to analyze the catch data and conduct length-dependent catch comparison and catch ratio analyses. Using the catch information (numbers and sizes of cod or saithe in each gillnet set deployment), we wanted to determine whether there was a significant difference in the catch efficiency averaged over deployments between the nylon PA gillnet and the biodegradable gillnet. We also wanted to determine if a potential difference between the gillnet types could be related to the size of the cod or saithe. The analysis was conducted separately for each species (cod and saithe) and each biodegradable gillnet (0.55 mm and 0.60 mm) following the procedure described below.

To assess the relative length-dependent catch efficiency effect of changing from nylon PA gillnet to a biodegradable gillnet, we used the method described in [29] and compared the catch data for the two net types. This method models the length-dependent catch comparison rate (*CC*_*l*_) summed over gillnet set deployments (for the full deployment period):

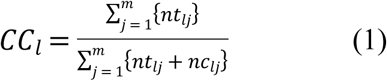

where *nc*_*lj*_ and *nt*_*lj*_ are the numbers of cod caught in each length class *l* for the nylon PA gillnet (control) and the biodegradable gillnet (treatment) in deployment *j* of a gillnet set (first or second set). *m* is the number of deployments carried out with one of the two sets. The functional form for the catch comparison rate *CC(l*,***v***) (the experimental being expressed by equation 1) was obtained using maximum likelihood estimation by minimizing the following expression:

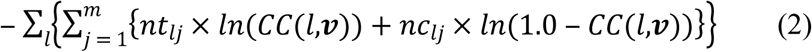

where ***v*** is a vector of the parameters describing the catch comparison curve defined by *CC(l*,***v***). The outer summation in the equation is the summation over length classes *l*. When the catch efficiency of the biodegradable gillnet and nylon PA gillnet is similar, the expected value for the summed catch comparison rate would be 0.5. Therefore, this baseline can be applied to judge whether or not there is a difference in catch efficiency between the two gillnet types. The experimental *CCl* was modelled by the function *CC(l*,***v***) using the following equation:

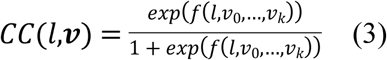

where *f* is a polynomial of order *k* with coefficients *v*_*0*_ to *v*_*k*_. The values of the parameters ***v*** describing *CC(l*,***v***) were estimated by minimizing equation (2), which was equivalent to maximizing the likelihood of the observed catch data. We considered *f* of up to an order of 4 with parameters *v*_*0*_, *v*_*1*_, *v*_*2*_, *v*_*3*_, and *v*_*4*_. Leaving out one or more of the parameters *v*_*0*_…*v*_*4*_ led to 31 additional models that were also considered as potential models for the catch comparison *CC(l*,***v***). Among these models, estimations of the catch comparison rate were made using multi-model inference to obtain a combined model [29, 30].

The ability of the combined model to describe the experimental data was evaluated based on the p-value. The p-value, which was calculated based on the model deviance and the degrees of freedom, should not be < 0.05 for the combined model to describe the experimental data sufficiently well, except for cases where the data are subject to over-dispersion [29, 31]. Based on the estimated catch comparison function *CC(l*,***v***) we obtained the relative catch efficiency (also named catch ratio) *CR(l*,***v***) between the two gillnet types using the following relationship:

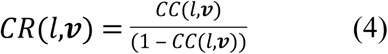

The catch ratio is a value that represents the relationship between catch efficiency of the biodegradable gillnet and that of the nylon PA gillnet. Thus, if the catch efficiency of both gillnets is equal, *CR(l*,***v***) should always be 1.0. *CR(l*,***v***) = 1.5 would mean that the biodegradable gillnet is catching 50% more cod of length l than the nylon PA gillnet. In contrast, *CR(l*,***v***) = 0.8 would mean that the biodegradable gillnet is only catching 80% of the cod of length l that the nylon PA gillnet is catching.

The confidence limits for the catch comparison curve and catch ratio curve were estimated using a double bootstrapping method [29]. This bootstrapping method accounts for between-set variability (the uncertainty in the estimation resulting from set deployment variation of catch efficiency in the gillnets and in the availability of cod) as well as within-set variability (uncertainty about the size structure of the catch for the individual deployments). However, contrary to the double bootstrapping method [29], the outer bootstrapping loop in the current study, which accounts for between deployment variation, was performed as a paired analysis for the biodegradable gillnet and nylon PA gillnet, taking full advantage of the experimental design with both nets being deployed simultaneously (see Fig. 1). By multi-model inference in each bootstrap iteration, the method also accounted for the uncertainty due to uncertainty in model selection. We performed 1,000 bootstrap repetitions and calculated the Efron 95% [32] confidence limits. To identify sizes of cod with significant differences in catch efficiency, we checked for length classes in which the 95% confidence limits for the catch ratio curve did not contain 1.0.

**Fig 1:**
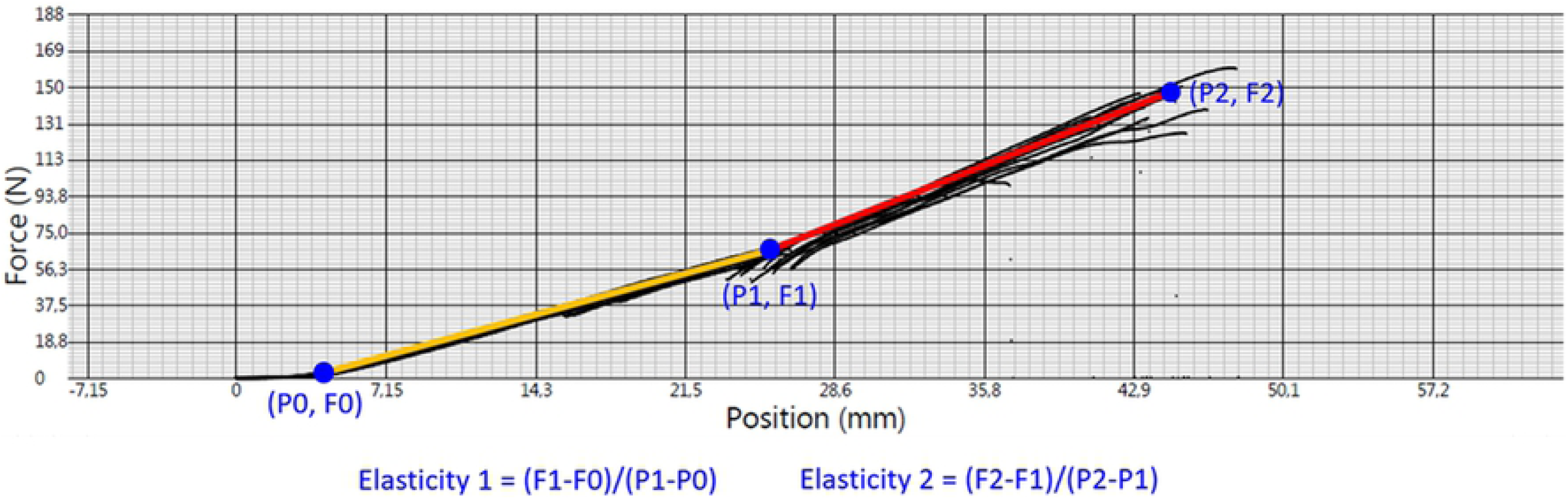
Elasticity. Estimation of *E*_*1*_ and *E*_*2*_ from force–displacement curve.

Finally, a length-integrated average value for the catch ratio was estimated directly from the experimental catch data using the following equation:

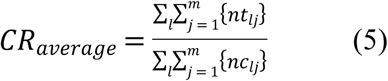

where the outer summation covers the length classes in the catch during the experimental fishing period.

#### Assessing the catch ratio of the two biodegradable gillnet designs

Because the same nylon gillnet design was used as a baseline in the asssessment of the catch ratio curves for both the 0.55 and 0.60 mm biodegradable gillnet, it was possible to indirectly assess the catch ratio curve between the two biodegradable gillnets. This was performed by calculating the ratio between the catch ratio curves obtained from the two catch ratio curves against the nylon net using the following equation:

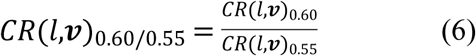

The 95% confidence intervals for *CR*(*l,v)*_0.60/0.55_were obtained based on the two the two bootstrap populations of results (1000 bootstrap repetitions in each) from each CR curve estimated for the 0.55 and 0.60 mm biodegradable gillnets against the nylon net. Since both bootstrap populations were obtained independently and the sampling to obtain those populations of results was performed randomly and independently, a new population of results with 1000 bootstrap iterations was created for *CR*(*l,v)*_0.60/0.55_following [33, 34]:

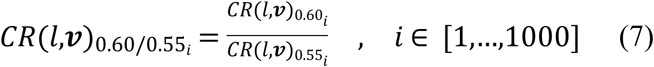

Where *i* represents the bootstrap repetion index. Based on this new population the Efron 95% confidence bands for *CR*(*l,v)*_0.60/0.55_ were obtained.

#### Assessment of mechanical properties

We estimated the mean tensile strength, elongation at break and the elasticity of the samples. Tensile strength, defined as the stress needed to break the sample, is given in kg. Elongation at break, defined as the length of the sample after it had stretched to the point when it breaks, is given as a percentage relative to the initial mesh size. Elasticity is a measurement of the resistance of an object or substance to being deformed elastically (stiffness) when a force is applied to it. The outputs from tensile testing were force-displacement curves which are described by the followeing equation:

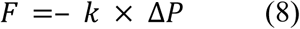

where Δ*P* is the amount of deformation (displacement in mm) produced by the force *F*, and *k* is a proportionality constant that depends on the shape and composition of the object and the direction of the force. We estimated two elasticities from the slopes of the force–displacement curve in the elastic deformation region (Fig 1).

Force-elongation curves were obtained from tensile strength testing for all types of gillnets, new and used. For each replicate, the tensile strength was determined as the peak of the force-elongation curve, and the corresponding elongation was taken as the elongation at break. For a set of samples, the tensile strength was determined as the average of all replicates, and polynomial fitting was performed to determine the average force-elongation curve.

Force-elongation curves of new and used gillnets from experiments carried out in 2017-2019 are presented and used in the discussion section to support the findings of this study.

## Results

Sufficient data was collected for both cod and saithe throughout the trial period. A total of 1,200 cod were caught, 780 using the nylon PA gillnet and 420 in the biodegradable gillnet (269 with the 0.55 mm and 151 with the 0.60 mm nets). A total of 1,328 saithe individuals were collected, of these, 736 were caught in the nylon PA gillnets and the remaining 592 were caught in the biodegradable gillnet (403 with the 0.55 mm and 189 with the 0.60 mm nets). Data were collected for 21 gillnet deployments for both cod and saithe, but the analysis was conducted based on deployments that had at least 10 fish in each set (Table 1). This was done in order to reduce the potential for additional uncertainty in the results and has been used successfully in previous catch comparison studies [18, 19].

**Table 1:**
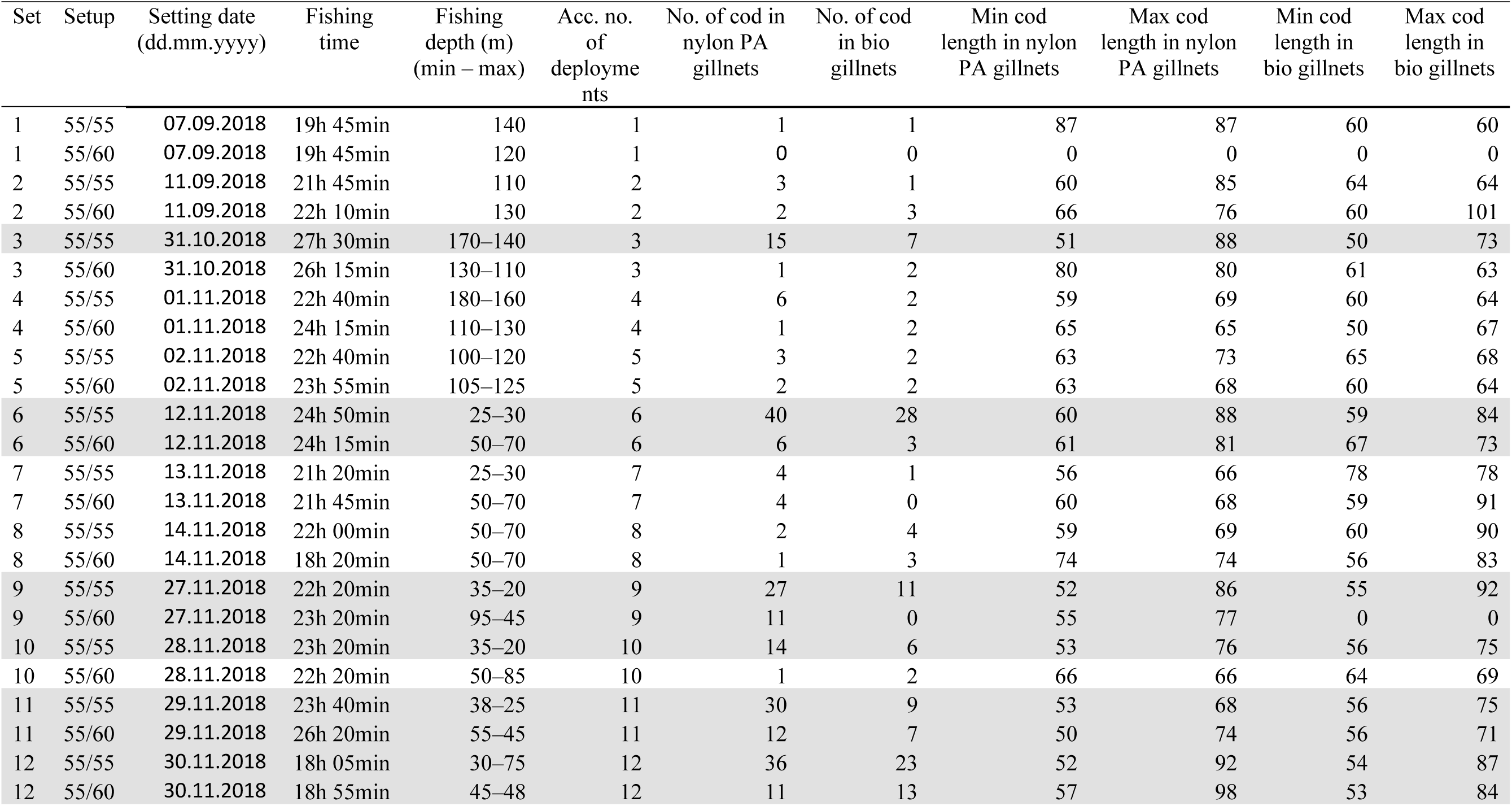
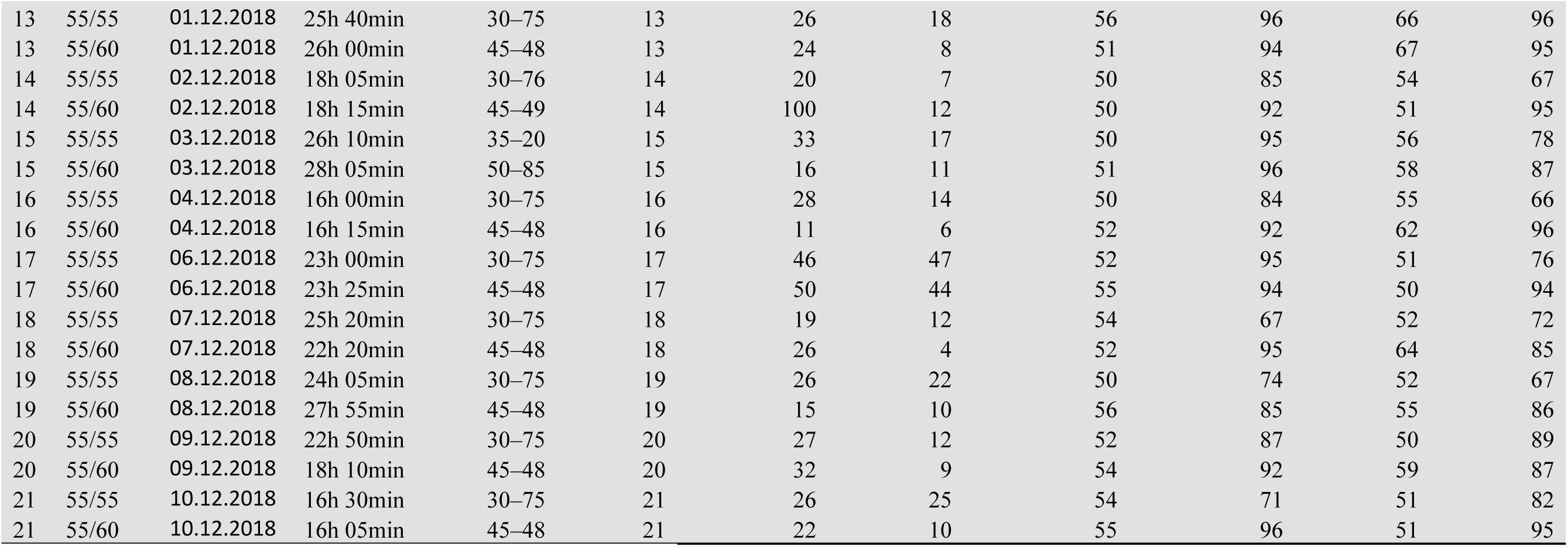
Catch data from all deployments for cod. The rows highlighted in grey indicate sets used in the analysis (sets containing at least 10 cod). The setups with 55 mm nylon PA gillnets / 55 or 60 mm biodegradable gillnets are indicated by 55/55 and 55/60.

| Set | Setup | Setting date<br>(dd.mm.yyyy) | Fishing<br>time | Fishing<br>depth (m)<br>(min – max) | Acc. no.<br>of<br>deployments | No. of cod in<br>nylon PA<br>gillnets | No. of cod<br>in bio<br>gillnets | Min cod<br>length in nylon<br>PA gillnets | Max cod<br>length in nylon<br>PA gillnets | Min cod<br>length in<br>bio gillnets | Max cod<br>length in<br>bio gillnets |
| --- | --- | --- | --- | --- | --- | --- | --- | --- | --- | --- | --- |
| 1 | 55/55 | 07.09.2018 | 19h 45min | 140 | 1 | 1 | 1 | 87 | 87 | 60 | 60 |
| 1 | 55/60 | 07.09.2018 | 19h 45min | 120 | 1 | 0 | 0 | 0 | 0 | 0 | 0 |
| 2 | 55/55 | 11.09.2018 | 21h 45min | 110 | 2 | 3 | 1 | 60 | 85 | 64 | 64 |
| 2 | 55/60 | 11.09.2018 | 22h 10min | 130 | 2 | 2 | 3 | 66 | 76 | 60 | 101 |
| 3 | 55/55 | 31.10.2018 | 27h 30min | 170–140 | 3 | 15 | 7 | 51 | 88 | 50 | 73 |
| 3 | 55/60 | 31.10.2018 | 26h 15min | 130–110 | 3 | 1 | 2 | 80 | 80 | 61 | 63 |
| 4 | 55/55 | 01.11.2018 | 22h 40min | 180–160 | 4 | 6 | 2 | 59 | 69 | 60 | 64 |
| 4 | 55/60 | 01.11.2018 | 24h 15min | 110–130 | 4 | 1 | 2 | 65 | 65 | 50 | 67 |
| 5 | 55/55 | 02.11.2018 | 22h 40min | 100–120 | 5 | 3 | 2 | 63 | 73 | 65 | 68 |
| 5 | 55/60 | 02.11.2018 | 23h 55min | 105–125 | 5 | 2 | 2 | 63 | 68 | 60 | 64 |
| 6 | 55/55 | 12.11.2018 | 24h 50min | 25–30 | 6 | 40 | 28 | 60 | 88 | 59 | 84 |
| 6 | 55/60 | 12.11.2018 | 24h 15min | 50–70 | 6 | 6 | 3 | 61 | 81 | 67 | 73 |
| 7 | 55/55 | 13.11.2018 | 21h 20min | 25–30 | 7 | 4 | 1 | 56 | 66 | 78 | 78 |
| 7 | 55/60 | 13.11.2018 | 21h 45min | 50–70 | 7 | 4 | 0 | 60 | 68 | 59 | 91 |
| 8 | 55/55 | 14.11.2018 | 22h 00min | 50–70 | 8 | 2 | 4 | 59 | 69 | 60 | 90 |
| 8 | 55/60 | 14.11.2018 | 18h 20min | 50–70 | 8 | 1 | 3 | 74 | 74 | 56 | 83 |
| 9 | 55/55 | 27.11.2018 | 22h 20min | 35–20 | 9 | 27 | 11 | 52 | 86 | 55 | 92 |
| 9 | 55/60 | 27.11.2018 | 23h 20min | 95–45 | 9 | 11 | 0 | 55 | 77 | 0 | 0 |
| 10 | 55/55 | 28.11.2018 | 23h 20min | 35–20 | 10 | 14 | 6 | 53 | 76 | 56 | 75 |
| 10 | 55/60 | 28.11.2018 | 22h 20min | 50–85 | 10 | 1 | 2 | 66 | 66 | 64 | 69 |
| 11 | 55/55 | 29.11.2018 | 23h 40min | 38–25 | 11 | 30 | 9 | 53 | 68 | 56 | 75 |
| 11 | 55/60 | 29.11.2018 | 26h 20min | 55–45 | 11 | 12 | 7 | 50 | 74 | 56 | 71 |
| 12 | 55/55 | 30.11.2018 | 18h 05min | 30–75 | 12 | 36 | 23 | 52 | 92 | 54 | 87 |
| 12 | 55/60 | 30.11.2018 | 18h 55min | 45–48 | 12 | 11 | 13 | 57 | 98 | 53 | 84 |
| 13 | 55/55 | 01.12.2018 | 25h 40min | 30–75 | 13 | 26 | 18 | 56 | 96 | 66 | 96 |
| 13 | 55/60 | 01.12.2018 | 26h 00min | 45–48 | 13 | 24 | 8 | 51 | 94 | 67 | 95 |
| 14 | 55/55 | 02.12.2018 | 18h 05min | 30–76 | 14 | 20 | 7 | 50 | 85 | 54 | 67 |
| 14 | 55/60 | 02.12.2018 | 18h 15min | 45–49 | 14 | 100 | 12 | 50 | 92 | 51 | 95 |
| 15 | 55/55 | 03.12.2018 | 26h 10min | 35–20 | 15 | 33 | 17 | 50 | 95 | 56 | 78 |
| 15 | 55/60 | 03.12.2018 | 28h 05min | 50–85 | 15 | 16 | 11 | 51 | 96 | 58 | 87 |
| 16 | 55/55 | 04.12.2018 | 16h 00min | 30–75 | 16 | 28 | 14 | 50 | 84 | 55 | 66 |
| 16 | 55/60 | 04.12.2018 | 16h 15min | 45–48 | 16 | 11 | 6 | 52 | 92 | 62 | 96 |
| 17 | 55/55 | 06.12.2018 | 23h 00min | 30–75 | 17 | 46 | 47 | 52 | 95 | 51 | 76 |
| 17 | 55/60 | 06.12.2018 | 23h 25min | 45–48 | 17 | 50 | 44 | 55 | 94 | 50 | 94 |
| 18 | 55/55 | 07.12.2018 | 25h 20min | 30–75 | 18 | 19 | 12 | 54 | 67 | 52 | 72 |
| 18 | 55/60 | 07.12.2018 | 22h 20min | 45–48 | 18 | 26 | 4 | 52 | 95 | 64 | 85 |
| 19 | 55/55 | 08.12.2018 | 24h 05min | 30–75 | 19 | 26 | 22 | 50 | 74 | 52 | 67 |
| 19 | 55/60 | 08.12.2018 | 27h 55min | 45–48 | 19 | 15 | 10 | 56 | 85 | 55 | 86 |
| 20 | 55/55 | 09.12.2018 | 22h 50min | 30–75 | 20 | 27 | 12 | 52 | 87 | 50 | 89 |
| 20 | 55/60 | 09.12.2018 | 18h 10min | 45–48 | 20 | 32 | 9 | 54 | 92 | 59 | 87 |
| 21 | 55/55 | 10.12.2018 | 16h 30min | 30–75 | 21 | 26 | 25 | 54 | 71 | 51 | 82 |
| 21 | 55/60 | 10.12.2018 | 16h 05min | 45–48 | 21 | 22 | 10 | 55 | 96 | 51 | 95 |

### Cod

For cod, this resulted in a total of 15 sets for analysis from the 0.55 mm setup and 12 from the 0.60 mm setup (Table 1). In the case of cod, both biodegradable gillnets (0.55 and 0.60 mm) had a significantly lower catch efficiency compared to the traditional nylon PA gillnet (0.55 mm) with estimated efficiencies of 62.38% (CI: 50.55–74.04) and 54.96% (CI: 35.42–73.52) compared with the nylon PA net, respectively (Tables 2 and Fig 2).

**Table 2:** Catch rate and fit statistic results from the 0.55 and 0.60 mm biodegradable gillnets vs. the 0.55 mm nylon PA set based on valid deployments for cod. Values in parentheses represent 95% confidence intervals. DOF denotes degrees of freedom.

| Length (cm) | Catch ratio (%) |  |
| --- | --- | --- |
|  | 0.55 mm | 0.60 mm |
| 50 | 74.59 (24.39–269.67) | 65.93 (24.43–410.77) |
| 55 | 70.97 (46.14–96.63) | 58.57 (28.60–139.11) |
| 60 | 66.97 (47.25–87.92) | 54.41 (29.05–94.91) |
| 65 | 62.66 (47.73–84.43) | 52.63 (29.65–74.56) |
| 70 | 58.17 (40.29–82.65) | 52.61 (30.64–70.73) |
| 75 | 53.72 (29.74–80.38) | 53.90 (31.20–83.56) |
| 80 | 48.70 (21.37–70.54) | 55.90 (33.45–106.62) |
| 85 | 45.71 (13.67–72.52) | 57.63 (33.27–126.53) |
| 90 | 42.56 (4.97–93.69) | 57.74 (28.19–116.21) |
| 95 | 40.37 (1.62–320.05) | 55.26 (9.76–109.90) |
| Average | 62.38 (50.55–74.04) | 54.96 (35.42–73.52) |
| P-value | 0.2915 | 0.0334<sup>†</sup> |
| Deviance | 45.46 | 60.29 |
| DOF | 41 | 42 |

**Fig 2:**
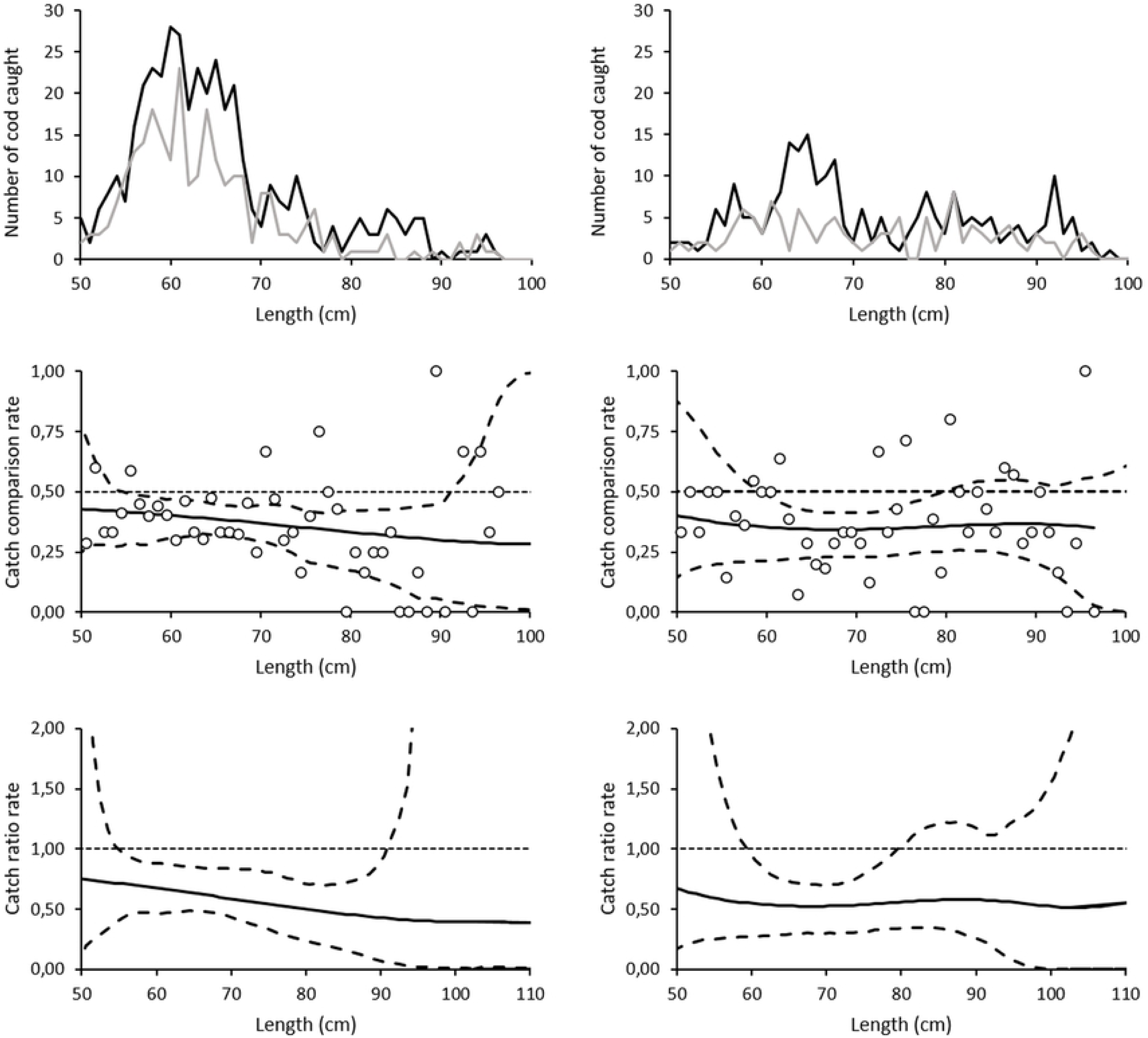
Size distribution, catch comparison rate and catch ratio rate for cod. The upper figures show the size distribution of cod caught using 0.55 mm nylon PA (black), and 0.55 mm (left) and 0.60 mm (right) biodegradable (grey) twine gillnets. The figures in the middle show the catch comparison curve for cod, with circle marks indicating the experimental rate, and the curve indicates the modelled catch comparison rate. The dotted line at 0.5 indicates the baseline where both gillnets fish the same amount. The dashed curves represent the 95% confidence interval for the estimated catch comparison curve. The lowest figure shows the estimated catch ratio curve for cod (solid line). The dotted line at 1.0 indicates the baseline where the fishing efficiency of both gillnet types is equal. The dashed curves represent the 95% confidence intervals of the estimated catch ratio curve.

Increasing the monofilament diameter from 0.55 mm to 0.60 mm did not have a significant effect on the catch efficiency of biodegradable gillnets. Both types of gillnets caught a similar number of cod in all length classes (Fig 3).

**Fig 3:**
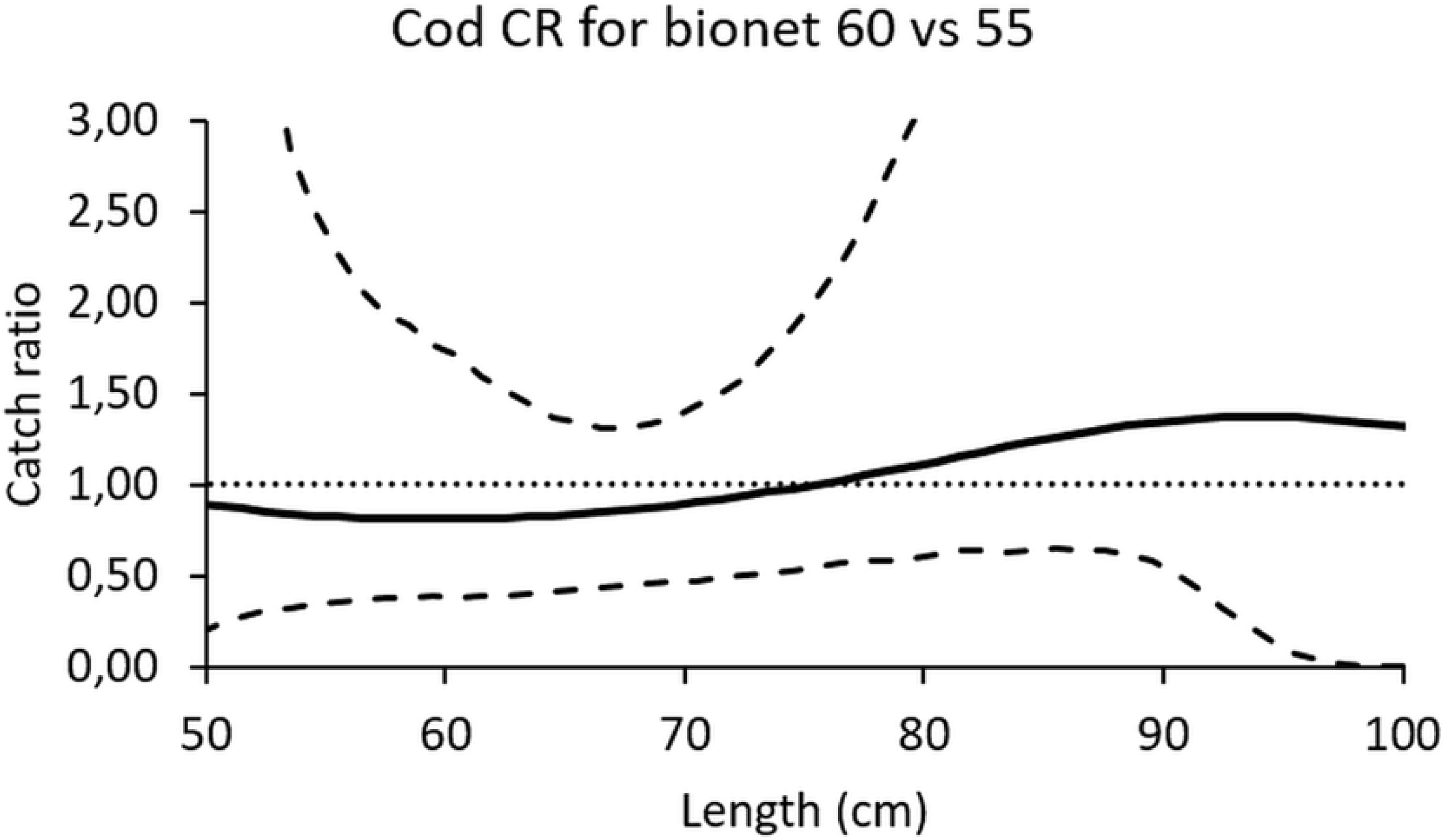
The estimated catch ratio curve for cod (solid line) The dashed curves represent the 95% confidence intervals of the estimated catch ratio curve. The dotted line at 1.0 indicates the baseline where the fishing efficiency of both gillnet types is equal.

### Saithe

For saithe, there were 15 sets for analysis of the 0.55 mm setup and 11 for the 0.60 mm setup (Table 4). Both biodegradable gillnets (0.55 and 0.60 mm) had a significantly lower catch efficiency for saithe compared to the traditional nylon PA net (0.55 mm) with estimated efficiencies of 83.40% (71.34–94.86) and 83.87% (66.36–104.92) compared with the nylon PA net, respectively (Table 5 and Fig 4).

**Table 4:** Catch data from all deployments for saithe. The rows highlighted in grey indicates sets used in the analysis (sets containing at least 10 saithe). The setups with 55 mm nylon PA gillnets / 55 or 60 mm biodegradable gillnets are indicated by 55/55 and 55/60.

| Set | Setup | Setting date<br>(dd.mm.yyyy) | Fishing<br>time | Fishing depth (m)<br>(min – max) | Acc. no. of<br>deployments | No. of saithe<br>in nylon PA<br>gillnets | No. of saithe<br>in bio<br>gillnets | Min. saithe<br>length in<br>nylon PA<br>gillnets | Max. saithe<br>length in<br>nylon PA<br>gillnets | Min. saithe<br>length in<br>bio gillnets | Max. saithe<br>length in<br>bio gillnets |
| --- | --- | --- | --- | --- | --- | --- | --- | --- | --- | --- | --- |
| 1 | 55/55 | 07.09.2018 | 19h 45min | 140 | 1 | 4 | 2 | 64 | 74 | 64 | 67 |
| 1 | 55/60 | 07.09.2018 | 19h 45min | 120 | 1 | 0 | 0 | 0 | 0 | 0 | 0 |
| 2 | 55/55 | 11.09.2018 | 21h 45min | 110 | 2 | 3 | 0 | 73 | 83 | 0 | 0 |
| 2 | 55/60 | 11.09.2018 | 22h 10min | 130 | 2 | 3 | 2 | 67 | 70 | 69 | 73 |
| 3 | 55/55 | 31.10.2018 | 27h 30min | 170–140 | 3 | 9 | 4 | 54 | 69 | 50 | 75 |
| 3 | 55/60 | 31.10.2018 | 26h 15min | 130–110 | 3 | 3 | 0 | 50 | 75 | 0 | 0 |
| 4 | 55/55 | 01.11.2018 | 22h 40min | 180–160 | 4 | 3 | 1 | 65 | 76 | 70 | 70 |
| 4 | 55/60 | 01.11.2018 | 24h 15min | 110–130 | 4 | 0 | 1 | 0 | 0 | 50 | 50 |
| 5 | 55/55 | 02.11.2018 | 22h 40min | 100–120 | 5 | 4 | 2 | 62 | 77 | 63 | 70 |
| 5 | 55/60 | 02.11.2018 | 23h 55min | 105–125 | 5 | 5 | 3 | 61 | 71 | 59 | 68 |
| 6 | 55/55 | 12.11.2018 | 24h 50min | 25–30 | 6 | 21 | 13 | 59 | 83 | 59 | 86 |
| 6 | 55/60 | 12.11.2018 | 24h 15min | 50–70 | 6 | 17 | 8 | 52 | 87 | 56 | 77 |
| 7 | 55/55 | 13.11.2018 | 21h 20min | 25–30 | 7 | 3 | 1 | 67 | 72 | 68 | 68 |
| 7 | 55/60 | 13.11.2018 | 21h 45min | 50–70 | 7 | 10 | 3 | 64 | 88 | 65 | 81 |
| 8 | 55/55 | 14.11.2018 | 22h 00min | 50–70 | 8 | 4 | 0 | 65 | 82 | 0 | 0 |
| 8 | 55/60 | 14.11.2018 | 18h 20min | 50–70 | 8 | 6 | 0 | 65 | 86 | 0 | 0 |
| 9 | 55/55 | 27.11.2018 | 22h 20min | 35–20 | 9 | 47 | 42 | 50 | 91 | 50 | 86 |
| 9 | 55/60 | 27.11.2018 | 23h 20min | 95–45 | 9 | 8 | 3 | 62 | 79 | 58 | 76 |
| 10 | 55/55 | 28.11.2018 | 23h 20min | 35–20 | 10 | 17 | 13 | 51 | 72 | 50 | 63 |
| 10 | 55/60 | 28.11.2018 | 22h 20min | 50–85 | 10 | 0 | 0 | 0 | 0 | 0 | 0 |
| 11 | 55/55 | 29.11.2018 | 23h 40min | 38–25 | 11 | 25 | 33 | 50 | 81 | 50 | 85 |
| 11 | 55/60 | 29.11.2018 | 26h 20min | 55–45 | 11 | 27 | 17 | 53 | 80 | 54 | 77 |
| 12 | 55/55 | 30.11.2018 | 18h 05min | 30–75 | 12 | 34 | 30 | 50 | 81 | 50 | 88 |
| 12 | 55/60 | 30.11.2018 | 18h 55min | 45–48 | 12 | 2 | 6 | 70 | 80 | 65 | 77 |
| 13 | 55/55 | 01.12.2018 | 25h 40min | 30–75 | 13 | 28 | 23 | 50 | 92 | 60 | 85 |
| 13 | 55/60 | 01.12.2018 | 26h 00min | 45–48 | 13 | 6 | 3 | 61 | 72 | 67 | 80 |
| 14 | 55/55 | 02.12.2018 | 18h 05min | 30–76 | 14 | 26 | 20 | 50 | 82 | 54 | 77 |
| 14 | 55/60 | 02.12.2018 | 18h 15min | 45–49 | 14 | 2 | 7 | 75 | 75 | 57 | 79 |
| 15 | 55/55 | 03.12.2018 | 26h 10min | 35–20 | 15 | 44 | 33 | 50 | 78 | 51 | 80 |
| 15 | 55/60 | 03.12.2018 | 28h 05min | 50–85 | 15 | 20 | 19 | 61 | 88 | 55 | 81 |
| 16 | 55/55 | 04.12.2018 | 16h 00min | 30–75 | 16 | 16 | 15 | 50 | 78 | 53 | 73 |
| 16 | 55/60 | 04.12.2018 | 16h 15min | 45–48 | 16 | 9 | 12 | 54 | 85 | 58 | 84 |
| 17 | 55/55 | 06.12.2018 | 23h 00min | 30–75 | 17 | 26 | 23 | 51 | 78 | 51 | 76 |
| 17 | 55/60 | 06.12.2018 | 23h 25min | 45–48 | 17 | 61 | 52 | 59 | 96 | 55 | 87 |
| 18 | 55/55 | 07.12.2018 | 25h 20min | 30–75 | 18 | 31 | 11 | 50 | 73 | 50 | 70 |
| 18 | 55/60 | 07.12.2018 | 22h 20min | 45–48 | 18 | 3 | 11 | 62 | 75 | 57 | 77 |
| 19 | 55/55 | 08.12.2018 | 24h 05min | 30–75 | 19 | 51 | 40 | 50 | 86 | 50 | 84 |
| 19 | 55/60 | 08.12.2018 | 27h 55min | 45–48 | 19 | 20 | 12 | 53 | 88 | 61 | 81 |
| 20 | 55/55 | 09.12.2018 | 22h 50min | 30–75 | 20 | 54 | 39 | 50 | 81 | 50 | 82 |
| 20 | 55/60 | 09.12.2018 | 18h 10min | 45–48 | 20 | 15 | 9 | 53 | 77 | 54 | 85 |
| 21 | 55/55 | 10.12.2018 | 16h 30min | 30–75 | 21 | 47 | 58 | 52 | 76 | 50 | 86 |
| 21 | 55/60 | 10.12.2018 | 16h 05min | 45–48 | 21 | 22 | 21 | 50 | 82 | 55 | 72 |

**Table 5:** Catch rate and fit statistic results from the 0.55 and 0.60 mm biodegradable gillnets vs. the 0.55 mm nylon PA set based on valid deployments for saithe. Values in parentheses represent the 95% confidence intervals. DOF denotes the degrees of freedom.

| Length (cm) | Catch ratio (%) |  |
| --- | --- | --- |
|  | 0.55 mm | 0.60 mm |
| 50 | 103.33 (64.00–199.22) | 126.66 (70.30–608.14) |
| 55 | 94.42 (73.90–140.63) | 124.11 (76.96–319.85) |
| 60 | 86.58 (70.16–110.11) | 110.00 (70.75–186.24) |
| 65 | 80.20 (63.52–92.19) | 93.93 (60.67–137.33) |
| 70 | 75.54 (53.68–88.66) | 79.96 (53.35–110.59) |
| 75 | 72.85 (46.76–95.12) | 68.32 (46.18–97.93) |
| 80 | 72.49 (47.52–119.27) | 57.43 (36.45–96.40) |
| 85 | 75.14 (43.22–261.02) | 45.23 (25.14–79.05) |
| 90 | 81.86 (31.08–1550.13) | 32.05 (8.66–67.15) |
| 95 | 93.83 (19.72–8043.05) | 23.18 (1.29–62.48) |
| Average | 83.40 (71.34–94.86) | 83.87 (66.36–104.92) |
| P-value | 0.6438 | 0.4114 |
| Deviance | 33.29 | 35.19 |
| DOF | 37 | 34 |

**Fig 4:**
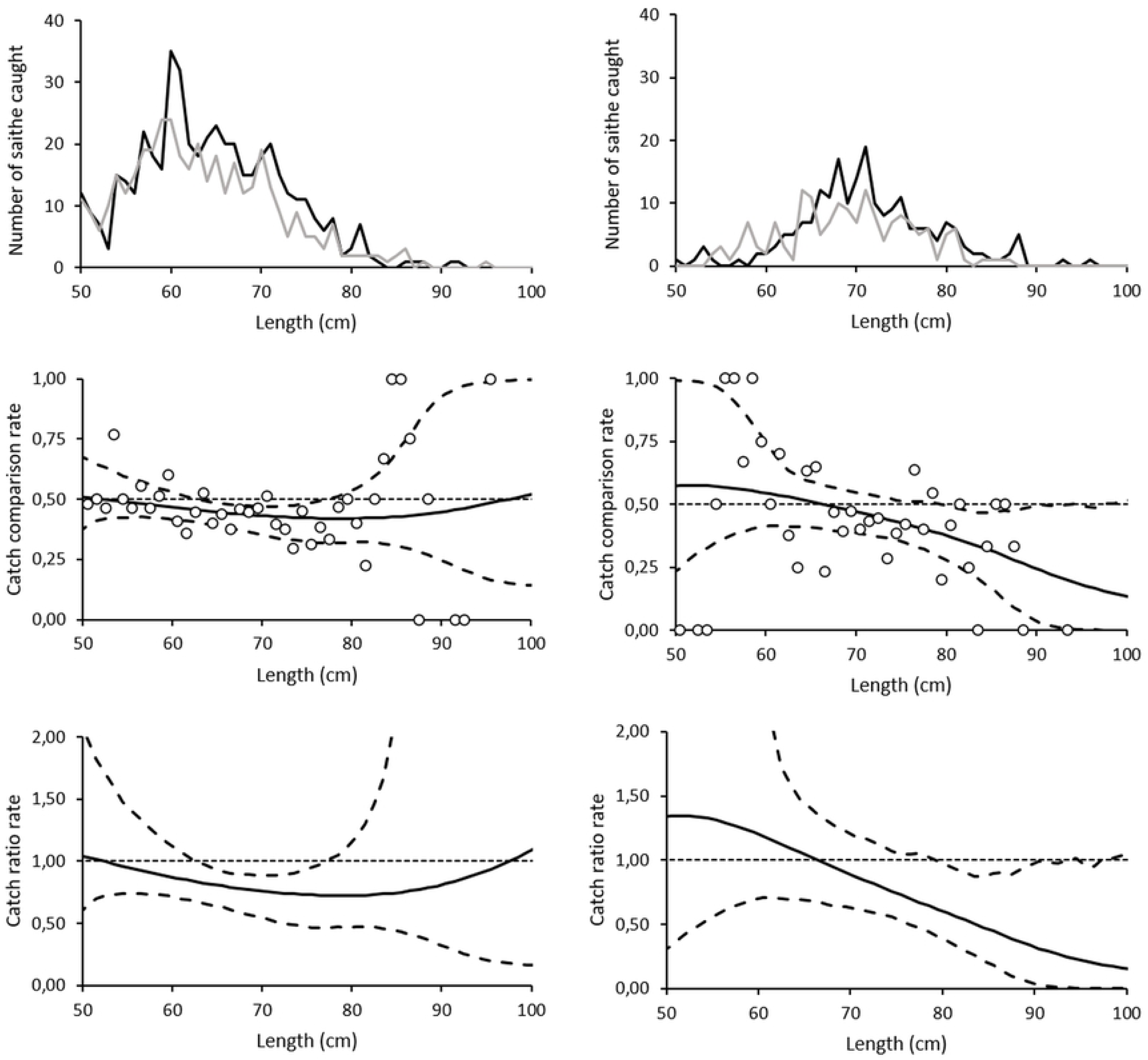
Size distribution, catch comparison rate and catch ratio rate for saithe. The upper figure shows the size distribution of saithe caught using the 0.55 mm nylon PA (black), and 0.55 mm (left) and 0.60mm (right) biodegradable (grey) twine gillnets. The figure in the middle shows the estimated catch ratio curve for saithe (solid line). The dotted line at 1.0 indicates the baseline where the fishing efficiency of both gillnet types is equal. The dashed curves represent the 95% confidence interval of the estimated catch ratio curve. The lowest figure shows the catch comparison curve for saithe, with circle marks indicating the experimental rate, and the curve indicates the modelled catch comparison rate. The dotted line at 1.0 indicates the baseline where fishing efficiency of both gillnet types is equal. The dashed curves represent the 95% confidence intervals of the estimated catch ratio curve.

Increasing the monofilament diameter from 0.55 mm to 0.60 mm did not have a significant effect on the catch efficiency of biodegradable gillnets. Both types of gillnets caught a similar number of saithe in all length classes (Fig 5).

**Fig 5:**
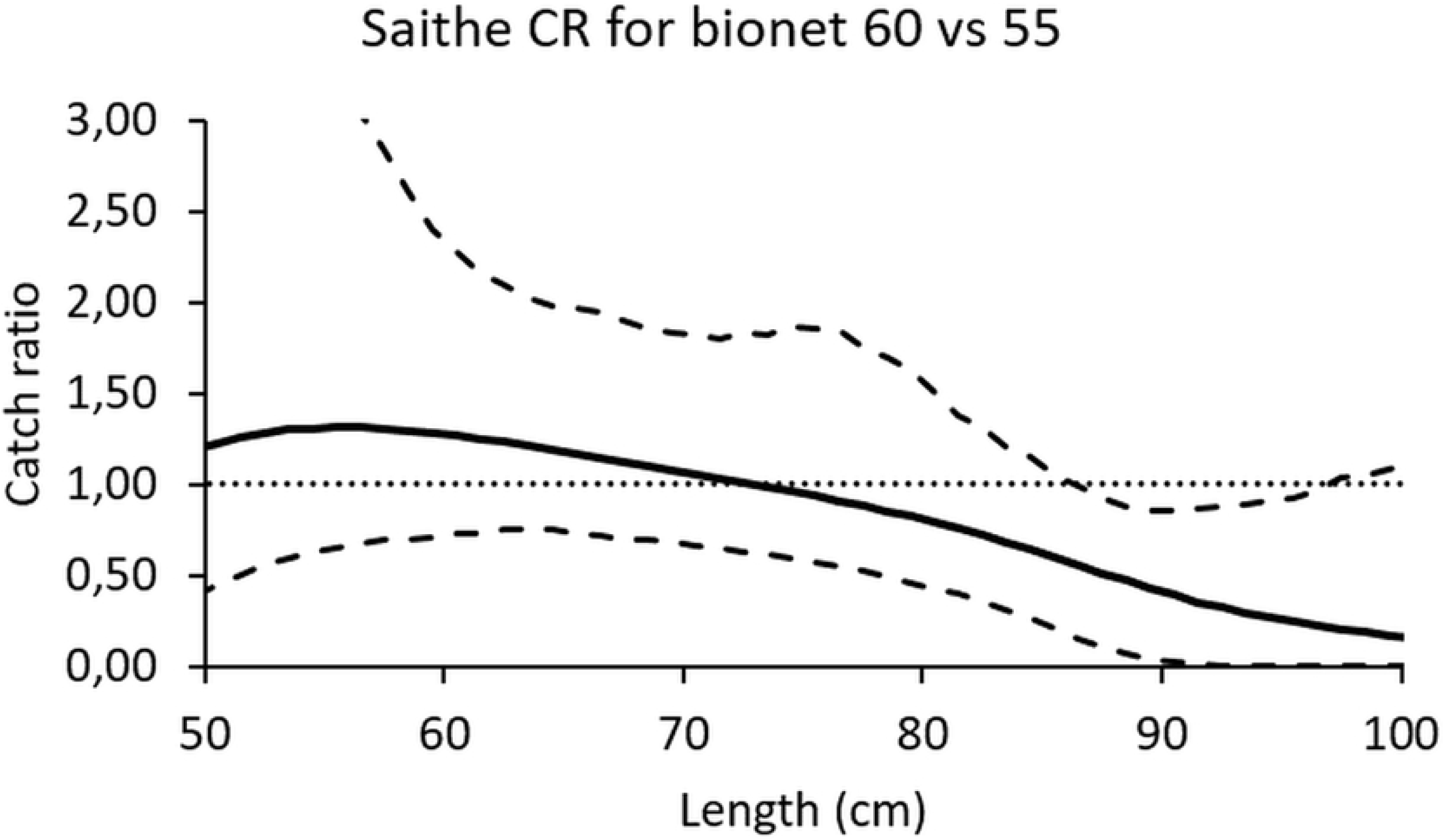
The estimated catch ratio curve for saithe (solid line) The dashed curves represent the 95% confidence interval of the estimated catch ratio curve. The dotted line at 1.0 indicates the baseline where fishing efficiency of both gillnet types is equal.

### Mechanical properties of the gillnets

New 0.55 mm nylon PA gillnets were 9.7% (t-test, p = 2.5×10^−5^) stronger than 0.55 mm biodegradable gillnets, and as strong as the 0.60 mm biodegradable gillnets (t-test, p = 0.402). New 0.55 mm nylon PA gillnets elongated significantly less at break than the 0.55 mm (17.0%; t-test, p = 7.1× 10^−17^) and 0.60 mm (16.6%; t-test, p = 1.6×10^−19^) biodegradable gillnets. The *E*_*1*_ and *E*_*2*_ of new nylon PA nets were significantly higher (t-test, p < 0.001) than the new 0.55 mm and 0.60 mm gillnets (Table 6).

**Table 6:**
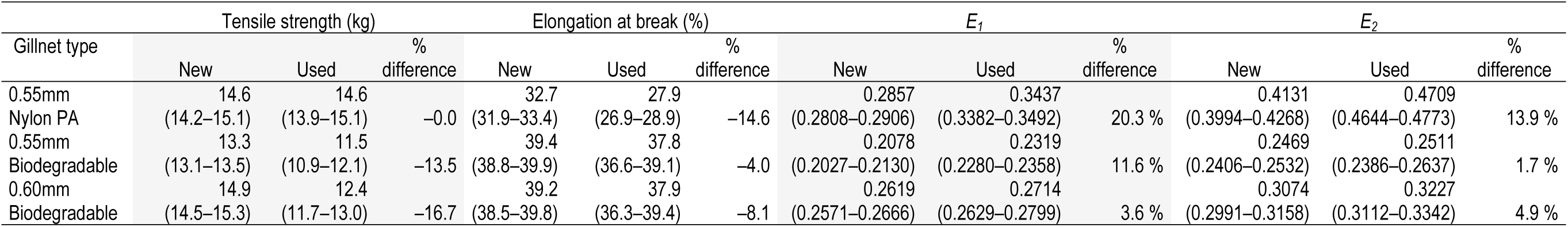
Mechanical properties of the gillnets. Mean tensile strength, elongation at break, *E*_*1*_ and *E*_*2*_ with 95% confidence intervals (in brackets) for new and used gillnets.

| Gillnet type | Tensile strength (kg) | | | Elongation at break (%) | | | $E_1$ | | | $E_2$ | | |
| --- | --- | --- | --- | --- | --- | --- | --- | --- | --- | --- | --- | --- |
|  | New | Used | % difference | New | Used | % difference | New | Used | % difference | New | Used | % difference |
| 0.55mm Nylon PA | 14.6<br>(14.2–15.1) | 14.6<br>(13.9–15.1) | –0.0 | 32.7<br>(31.9–33.4) | 27.9<br>(26.9–28.9) | –14.6 | 0.2857<br>(0.2808–0.2906) | 0.3437<br>(0.3382–0.3492) | 20.3 % | 0.4131<br>(0.3994–0.4268) | 0.4709<br>(0.4644–0.4773) | 13.9 % |
| 0.55mm Biodegradable | 13.3<br>(13.1–13.5) | 11.5<br>(10.9–12.1) | –13.5 | 39.4<br>(38.8–39.9) | 37.8<br>(36.6–39.1) | –4.0 | 0.2078<br>(0.2027–0.2130) | 0.2319<br>(0.2280–0.2358) | 11.6 % | 0.2469<br>(0.2406–0.2532) | 0.2511<br>(0.2386–0.2637) | 1.7 % |
| 0.60mm Biodegradable | 14.9<br>(14.5–15.3) | 12.4<br>(11.7–13.0) | –16.7 | 39.2<br>(38.5–39.8) | 37.9<br>(36.3–39.4) | –8.1 | 0.2619<br>(0.2571–0.2666) | 0.2714<br>(0.2629–0.2799) | 3.6 % | 0.3074<br>(0.2991–0.3158) | 0.3227<br>(0.3112–0.3342) | 4.9 % |

Used 0.55 mm nylon PA gillnets were significantly stronger (26.9%; t-test, p = 1.7× 10^−8^) and (17.7%; t-test, p = 2.2×10^−5^) than 0.55mm and 0.60mm biodegradable gillnets, respectively. Used 0.55 mm nylon PA gillnets elongated significantly less (26.2%; t-test, p = 4×10^−14^) and (26.4%; t-test, p = 8.2×10^−12^) at break than 0.55 mm and 0.60 mm used biodegradable gillnets, respectively. The *E*_*1*_ and *E*_*2*_ of used nylon PA gillnets was significantly higher (t-test, p < 0.001) than that for 0.55 mm and 0.60 mm used biodegradable gillnets, respectively (Table 6).

Nylon PA gillnets were as strong, elongated 14.6% less at break (from 32.7 to 27.9%; t-test, p = 1.49× 10^−8^), and were significantly more elastic (*E*_*1*_ = 20.3% and *E*_2_ = 13.9%; t-test, p < 0.001) after having been deployed 21 times at sea. Both types of biodegradable gillnets suffered significant reductions in tensile strength (t-test, p < 0.001). The 0.55 mm biodegradable gillnet decreased from 13.3 to 11.5 kg and the 0.60 mm biodegradable gillnet decreased from 14.9 to 12.4 kg after being used 21 times at sea. The 0.55 and 0.60 mm nets elongated significantly less at break (4.0%; t-test, p = 3.31× 10^−2^) and (8.1%; t-test, p = 7.70×10^−4^), respectively, and *E*_*1*_ and *E*_*2*_ increased after use (Table 6).

The fitted force-elongation curves from tensile testing (Fig 6) shows that used nylon PA gillnets exhibited an increase in stiffness, while used biodegradable gillnets experienced a slight decrease.

**Fig 6:**
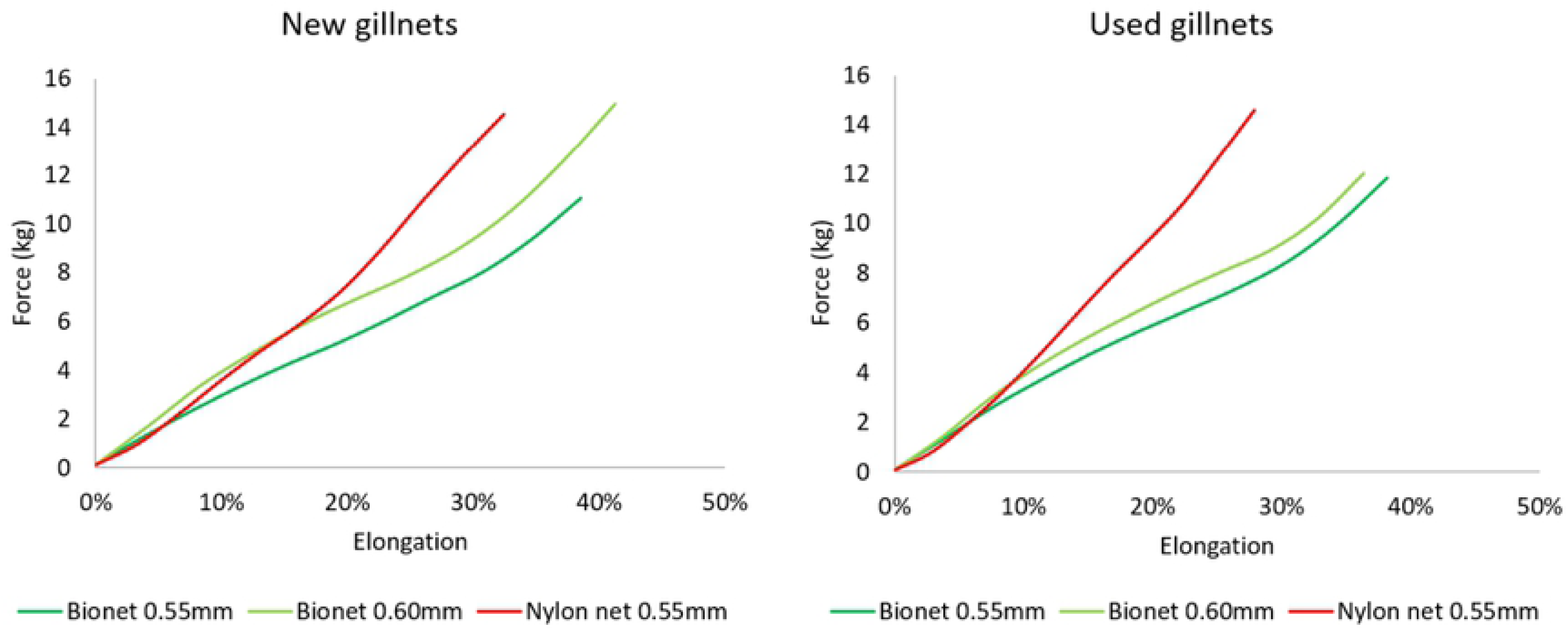
Force–elongation curves of new and used gillnets. Elongation is shown as a percentage relative to the initial length.

## Discussion

Increasing the monofilament thickness of biodegradable gillnets from 0.55 to 0.60 mm to match the tensile strength of the 0.55 mm nylon PA gillnets did not improve their catch efficiency. No difference in breaking strength between 0.55 mm nylon PA and 0.60 mm biodegradable gillnets was detected when the gillnets were new. However, the 0.55 mm nylon PA gillnets caught significantly more cod and saithe than the 0.60 mm biodegradable gillnets during the fishing season and generally showed better catch rates for most length classes. Our results are consistent with those reported by Grimaldo, et al. [18, 19] for the catch characteristics of gillnets for cod, saithe and Greenland halibut (*Reinhardtius hippoglossoides*), those of Bae et al. [24] for flounder (*Cleisthenes pinetorum*), and those of Kim et al. for yellow croaker (*Larimichthys polyactis*). These researchers found that the fishing efficiency of nylon PA gillnets was 1.1- to 1.4-times higher than biodegradable gillnets and concluded that differences in the mechanical properties of the materials (i.e., tensile strength) could explain the differences in catch efficiency. All of these studies showed that biodegradable gillnets were generally 10–16% weaker and elongate 8–10% more at break than nylon PA gillnets of similar twine diameter. However, none of these studies carried out a more comprehensive assessment of the potential effects of other mechanical properties (i.e., elongation, elasticity, stiffness) on the catch efficiency of the gillnets. The results of our study suggests that tensile strength may not be the main cause of the low catch efficiency of biodegradable gillnets relative to that of nylon PA gillnets, and we therefore speculate whether the elasticity and stiffness may better explain the catch efficiency patterns of nylon PA and biodegradable gillnets.

Significant differences in the elasticity and stiffness were found between biodegradable and nylon PA gillnets and therefore these two parameters may have caused the differences in catch efficiency between the gillnets. The increased stiffness of monofilaments can be identified as an increased slope in a force-elongation curve from tensile testing, and a change in the stiffness properties of the monofilaments after use may indicate degradation (or deterioration) of the polymer material. The fitted force-elongation curves from tensile testing shown in Fig 6 shows an increase in the stiffness of used nylon PA monofilaments, while the used biodegradable monofilaments experienced the opposite effect. The ratio of force-elongation is elasticity-stiffness, but only the force defines the strength of the material. Strength measures how much stress the material can handle before permanent deformation or fracture occurs, whereas stiffness measures the resistance to elastic deformation. In contrast to nylon PA gillnets, biodegradable gillnets increased in elasticity and reduced in stiffness after use. Based on these results, we speculate whether the biodegradable gillnets became too elastic and consequently fish could easily press themselves through the meshes of the gillnet and avoid capture. The force-elongation curves from earlier experiments obtained from biodegradable and nylon PA gillnet samples (Fig 7) give an indication of the differences in elongation and stiffness between these two types of gillnets. Although Fig 7 shows a large variation in the results for type of gillnets and year, it is possible to see a certain tendency for the nylon PA gillnets to be stiffer than the biodegradable gillnets, when new and used. It also seems that used biodegradable gillnets tend to become less stiff and elongate less than nylon PA gillnets after use.

**Fig 7:**
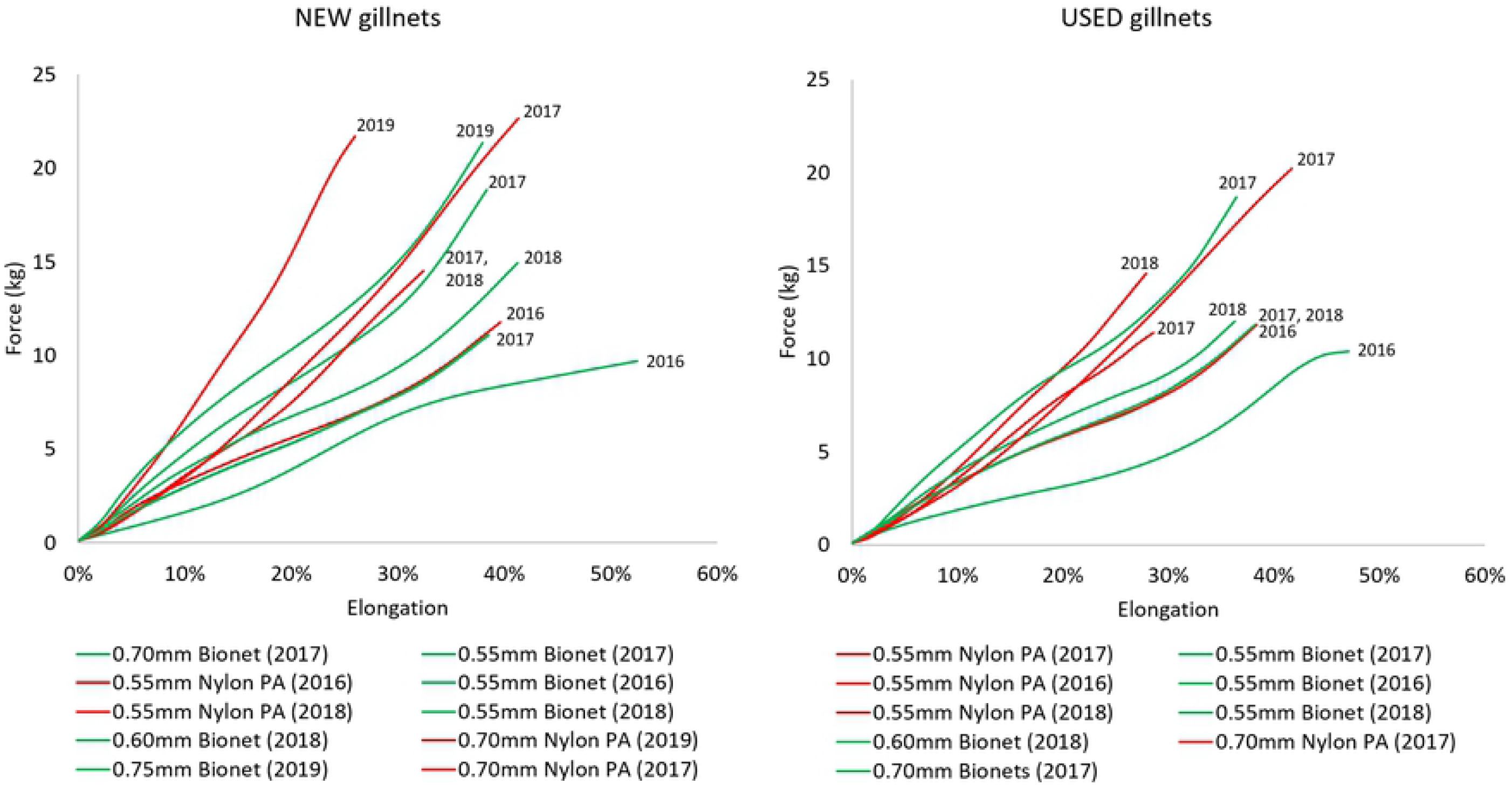
Force-elongation curves of new and used gillnets from experiments carried out in 2017-2019. Elongation is given as a percentage relative to the initial length.

The elasticity and stiffness of nylon PA and biodegradable materials are probably closely related to the way these two types of gillnet catch fish, better known as “catching modes” [35]. For instance, a stiffer and less elastic material may catch more fish by gilling, while a more flexible and elastic material can fish more by snagging. A quantification of the number and length distribution of fish caught per catching mode type can potentially provide information on the effect that elasticity and stiffness have on the catch efficiency of gillnets. This information can also be used for improving size selectivity and to narrow the wide selection range that traditional gillnets are known for. Knowing more about the effect of elasticity and stiffness on the caching modes can also lead to the enhancement of some catch methods to improve catch quality, since wedging and entangling are known to cause marks in the fish and reduce the quality of the filet, while snagging and gilling may yield better quality fish. Unfortunately, our experimental setups did not allow us to investigate how the elastic modulus affects the catch efficiency of the gillnets, and consequently this is only a hypothesis that should be investigated in future experiments.

The deterioration of nylon PA and biodegradable gillnets in this experiment was the result of chemical and mechanical changes that occurred during the three-month experimental period. Different mechanisms of degradation might have acted simultaneously on the nylon PA and biodegradable fibers, and some probably had a stronger effect than others. Although this experiment was unable to identify and quantify the effect of specific mechanisms of degradation of the gillnets that were studied, possible degradation mechanisms during the field experiments are microbiological degradation, hydrolysis, oxidation, and mechanical damage (i.e., abrasion in the hauling machine, friction due to contact with hard surfaces when the gillnets were operated on deck). Polymers are also known to also be vulnerable to UV-exposure, however since the experiment was carried out during the last part of the polar night period in northern Norway, we consider the effect of UV-radiation to be negligible.

## Acknowledgements

This study was financed by the Regional Research Council Region Northern Norway and the Norwegian Directorate of Fisheries. We are very grateful to captain Bent Gabrielsen and the crew on board the “MS Karoline” for their help in the fishing trials. We would like to thank S-ENPOL Co. Ltd. for their collaboration in this project and for providing the gillnet samples to carry out the experiments at sea.

## Supporting information

**S1 Fig. Catch data for individual sets for cod**. The catch data consists of count data for numbers of cod caught in the biodegradable gillnets (Test 1) and nylon PA gillnets (Test 2) for each size class (Length) corresponding to total fish length.

**S1 Fig. Catch data for individual sets for saithe**. The catch data consists of count data for numbers of saithe caught in the biodegradable gillnets (Test 1) and nylon PA gillnets (Test 2) for each size class (Length) corresponding to total fish length.

## References

[1] FAO. The State of World Fisheries and Aquaculture 2018 - Meeting the sustainable development goals. Rome. 2018. ISBN 978-92-5-130562-1.

[2] Fisheries Directorate of Norway (2020) Catches and licences https://www.fiskeridir.no/Yrkesfiske/Tall-og-analyse/Fangst-og-kvoter

[3] Fisheries Directorate of Norway (2020) Fishermen, vessels and permits. https://www.fiskeridir.no/Yrkesfiske/Tall-og-analyse/Fiskere-fartoey-og-tillatelser

[4] Deshpande PC, Philis G, Brattebø H, Fet AF. Using Material Flow Analysis (MFA) to generate the evidence on plastic waste management from commercial fishing gears in Norway. Resour conserve recy. 2020; 5: 100024. https://doi.org/10.1016/j.rcrx.2019.100024

[5] Sundt P, Briedis R, Skogesal O, Standal E, Rødås Johnsen H, Schulze P. Basis for assessing the producer responsibility scheme for the fishing and aquaculture industry. Report of the Norwegian Environmental Agency. 2018; M-1052, 178 pp.

[6] Macfadyen G, Huntington T, Cappell R. Abandoned, lost or otherwise discarded fishing gear. FAO Fisheries and Aquaculture Technical Paper, 2009 (No.523): p. 115 pp.

[7] Schaefer MB. Some Considerations of Population Dynamics and Economics in Relation to the Management of the Commercial Marine Fisheries. J Fish Res Board Can. 1957. 14(5): p. 669–681.

[8] Hareide NR, Garnes G, Rihan D, Mulligan M, Tyndall P, Clark M, et al. A Preliminary Investigation on Shelf Edge and Deepwater Fixed Net Fisheries to the West and North of Great Britain, Ireland, around Rockall and Hatton Bank. ICES CM 2005/ N:07, 2005.

[9] Large PA, Graham N, Hareide NR, Misund R, Rihan D, Mulligan M, et al. Lost and abandoned nets in deep-water gillnet fisheries in the Northeast Atlantic: retrieval exercises and outcomes. ICES J Mar Sci. 2009; 66(2): 323–333. https://doi.org/10.1093/icesjms/fsn220

[10] Norwegian Fisheries Directorate (2019). Lost fishing gears and ghost fishing. https://www.fiskeridir.no/Yrkesfiske/Areal-og-miljoe/Tapte-fiskeredskap

[11] Tokiwa Y, Calabia BP, Ugwu CU, Aiba S. Biodegradability of plastics. Int J Mol Sci. 2009, 10(9): 3722–3742. https://doi.org/10.3390/ijms10093722

[12] Park SW, Bae JH, Lim JH, Cha BJ, Park CD, Yang YS, Ahn HC. Development and physical properties on the monofilament for gill nets and traps using biodegradable aliphatic polybutylene succinate resin. J Kor Soc Fish Tech. 2007; 43(4): 281–290. https://doi.org/10.3796/KSFT.2007.43.4.281

[13] Park SW, Bae JH, Weatherability of biodegradable polybutylene succinate (PBS) monofilaments. J Kor Soc Fish Tech. 2008; 44(4): 265–272. https://doi.org/10.3796/KSFT.2008.44.4.265

[14] Kim S, Park S, Lee K, Lim J. Characteristics on the fishing performance of a drift net for yellow croaker (*Larimichthys polyactis*) in accordance with the thickness of a net twine. J Kor Soc Fish Tech. 2013; 49(3): 218–226. https://doi.org/10.3796/KSFT.2012.49.3.218

[15] Kim S, Park S, Lee K. Fishing performance of an Octopus minor net pot made of biodegradable twines. Turkish J Fish Aquat Sci. 2014a; 14: 21–30.

[16] Kim S, Park S, Lee K. Fishing performance of environmentally friendly tubular pots made of biodegradable resin (PBS/PBAT) for catching the conger eel Conger Myriaster. Fish Sci. 2014b; 80: 887–895. https://doi.org/10.1007/s12562-014-0785-z

[17] Kim S, Kim P, Lim J, An H, Suuronen P. Use of biodegradable driftnets to prevent ghost fishing: physical properties and fishing performance for yellow croaker. Anim Conserv. 2016; 19: 309–319. https://doi.org/10.1111/acv.12256

[18] Grimaldo E, Herrmann B, Tveit G, Vollstad J, Schei M. Effect of using biodegradable PBSAT gillnets on the catch efficiency and quality of Greenland halibut (*Reinhardtius hippoglossoides*). Mar Coast Fish. 2018a. 10:619–629. https://doi.org/10.1002/mcf2.10058|

[19] Grimaldo E, Herrmann B, Vollstad J, Su B, Moe Føre H, Larsen RB. Fishing efficiency of biodegradable PBSAT gillnets and conventional nylon gillnets used in Norwegian cod (*Gadus morhua*) and saithe (*Pollachius virens*) fisheries. ICES J Mar Sci. 2018b; 75(6): 2245–2256. https://doi.org/10.1093/icesjms/fsy108

[20] Grimaldo E, Herrmann, Vollstad J, Su B, Moe Føre H, Larsen RB. Comparison of fishing efficiency between biodegradable gillnets and conventional nylon gillnets. Fish Res. 2019; 213: 67–74. https://doi.org/10.1016/j.fishres.2019.01.003

[21] Urbanek AK, Rymowicz W, Strzelectki M, Kociuba W, Franczak K, Mironczuc A. Isolation and characterization of Arctic microorganisms decomposing bioplastics. AMB Express. 2017; 7(1): 148–148. https://doi.org/10.1186/s13568-017-0448-4

[22] Sekiguchi T, Saika A, Nomura K, Watanabe T, Fujimoto Y, Enoko M, et al. Biodegradation of aliphatic polyesters soaked in deep seawaters and isolation of poly(ε-caprolactone)-degrading bacteria. Polym Degrad Stab. 2011; 96(7): 1397–1403. https://doi.org/10.1016/j.polymdegradstab.2011.03.004

[23] Bae BS, Cho SK, Park SW, Kim, SH. Catch characteristics of the biodegradable gill net for flounder. J Kor Soc Fish Tech. 2012; 48(4): 310–321. https://doi.org/10.3796/KSFT.2012.48.4.310

[24] Bae BS, Lim JH, Park SW, Kim SH, Cho SK. Catch characteristics of gillnets for flounder by the physical properties of net filament in the East Sea. J Kor Soc Fish Tech. 2013; 49(2): 095–105. https://doi.org/10.3796/KSFT.2013.49.2.095

[25] Kim MK, Yun KC, Kang GD, Ahn JS, Kang SM, Kim YJ, et al. Biodegradable resin composition and fishing net produced from same. US 2017/0112111A1, 2017.

[26] Su B, Føre HM, Grimaldo E. A comparative study of the mechanical properties of biodegradable PBSAT and PA gillnets in Norwegian coastal waters. OMAE2019-95350 2019.

[27] Herrmann B, Sistiaga M, Nielsen KN, Larsen RB. Understanding the size selectivity of redfish (*Sebastes spp*.) in North Atlantic trawl codends. J Northw Atl Fish Sci. 2012; 44: 1–13. http://dx.doi.org/10.2960/J.v44.m680

[28] Herrmann B, Krag LA, Feekings J, Noack T. Understanding and predicting size selection in diamond-mesh cod ends for Danish seining: a study based on sea trials and computer simulations. Mar Coast Fish. 2016; 8: 277–291. https://doi.org/10.1080/19425120.2016.1161682

[29] Herrmann B, Sistiaga M, Rindahl L, Tatone I. Estimation of the effect of gear design changes on catch efficiency: Methodology and a case study for a Spanish longline fishery targeting hake (*Merluccius merluccius*). Fish Res. 2017; 185: 153–160. https://doi.org/10.1016/j.fishres.2016.09.013

[30] Burnham KP, Anderson DR. Model Selection and Multimodel Inference: A Practical Information-theoretic Approach. 2nd ed. Springer, New York. 2002. ISBN 978-0-387-22456-5.

[31] Wileman DA, Ferro RST, Fonteyne R, Millar RB. Manual of Methods of Measuring the Selectivity of Towed Fishing Gears. ICES Coop. Res. Rep. No. 215, 1996. ICES, Copenhagen, Denmark. ISSN 1017–6195.

[32] Efron B. The jackknife, the bootstrap and other resampling plans. In: SIAM Monograph No. 38, CBSM-NSF Regional Conference Series in Applied Mathematics, Philadelphia. 1982. ISBN: 978-0-89871-179-0.

[33] Herrmann B, Krag LA, Krafft BA. Size selection of Antarctic krill (*Euphausia superba*) in a commercial codend and trawl body. Fish Res. 2018; 207: 49–54.

[34] Veiga-Malta T, Feekings J, Herrmann B, Krag LA. Industry-led fishing gear development: Can it facilitate the process? Ocean Coast Manag. 2019; 177: 148–155. https://doi.org/10.1016/j.ocecoaman.2019.05.009

[35] Grati F, Bolognini L, Domenichetti F, Fabi G, Polidori P, Santelli A, et al. The effect of monofilament thickness on the catches of gillnets for common sole in the Mediterranean small-scale fishery. Fish Res. 2015. 164: 170–177. https://doi.org/10.1016/j.fishres.2014.11.014

